# The Coevolution of Promoters and Transcription Factors in Animal and Plant Cells

**DOI:** 10.1101/2020.04.20.050187

**Authors:** Jingsong Zhang, Xiangtian Yu, Zhixi Su, Shutao He, Yiwei Zhou, Hao Dai, Xiaohu Hao, Tao Zeng, Wen Wang, Luonan Chen

## Abstract

Coevolution has been acknowledged to play a significant role in driving the evolution of molecules and species. Promoters and transcription factors (TFs), especially their interactions, are key determinants for the regulation of gene expression. However, the evolutionary processes and mechanisms of promoter and TF interactions are still poorly understood. Here we conduct extensive physicochemical analyses of multi-omics sequences in 440 animal species and 223 plant species which span nearly one billion years of phylogeny. We demonstrate that promoters and TFs obey antagonistic coevolution in the animal kingdom while follow mutualistic coevolution in the plant kingdom. Furthermore, we reveal that such two coevolutionary strategies result in different evolutionary transitions of transcriptional networks in the two kingdoms. These results suggest that the two distinct coevolutionary mechanisms are likely to be major drivers of far greater genetic divergence between animals and plants, and open a new door to understanding the roles of promoters and TFs in tumor initiation and progression, and human ageing as well in molecular interactions and evolution.

## INTRODUCTION

Evolutionary interaction was first introduced by Charles Darwin to explain the interactions between flowering plants and insects. Coevolutionary theories are now well developed and have been applied in many fields, such as cosmology (Heckman and Kauffmann, 2011; Martin-Navarro et al., 2018; Wu et al., 2015). Sociology (Henrich et al., 2006), computer science (Liu and Lu, 2015), biology (Bottery et al., 2019; Chen et al., 2020; Cross et al., 2019; Garcia-Bayona and Comstock, 2018), biomedicine (Bottery et al., 2017; Loftie-Eaton et al., 2016; Single et al., 2007), and molecular biology (Barreto et al., 2018; Jin et al., 2017b; Lord et al., 2019; Mann et al., 2016). The interactions between promoters and transcription factors (TFs) are the beginning of gene transcription, and it has been acknowledged that such interactions are vital for transcription regulation and could affect all stages of cell lifecycle (Mirny, 2010). The evolution of interactions between promoters and TFs (Thomas and Chiang), along with eukaryotes, has continued for approximately 1.6 to 2.1 billion years (Bengtson et al., 2017; Zhu et al., 2016). Whether the interactions are coevolution, and if yes, what kind of coevolutionary relationships they follow, are key questions in molecular biology. Elucidation of these questions could offer a better understanding of biological evolution and ecological processes. However, existing studies on promoter-TF interactions are usually limited to the differences of cell types, tissues or developmental stages (Briggs et al., 1986; Cordingley et al., 1987; Whalen et al., 2016). In this study, we elucidated the evolutionary trajectories of promoter-TF interactions through nearly one billion years of evolutionary history by analyzing DNA and protein sequences.

Promoter stability is of primary importance for the initiation of gene transcription. Double-stranded breaks in DNA, especially in promoter regions, pose major threats to genome integrity because they may result in chromosomal aberrations (van Gent et al., 2001). GC-content (see STAR Methods) of promoters has a substantial impact on DNA molecular stability as well as gene activity, because the connection between G and C bases (three hydrogen bonds, G≡C) is stronger than that between A and T bases (two hydrogen bonds, A=T) and also there is more favorable stacking energy for GC pairs than for AT pairs (Yakovchuk et al., 2006). Actually, it is considered to be more difficult to open the double DNA strands of a gene by TFs if the GC-content on its promoter is higher. GC-content is found to be variable within a genome (Furey and Haussler, 2003) and among different organisms (Birdsell, 2002). Thus, it is desirable that we quest the evolution of promoters by studying the GC-content. In addition to GC-content, the isoelectric-point (pI) of molecules/residues is a crucial physicochemical property and a main biochemical criterion (Dika et al., 2015; Liu et al., 2009), which influence the structure and functions of proteins (including TFs). As such, we deem it is necessary to explore the coevolutionary trajectories of TFs and their corresponding promoters in the entire phylogeny by evaluating the variations of TF pIs and GC-content.

In this study, the benchmark datasets we used include both genome and proteome sequences (from AnimalTFDB (Hu et al., 2019), PlantTFDB (Jin et al., 2017a), Ensemble (Hunt et al., 2018), EnsemblePlants (Kersey et al., 2018) and UniProt (Bateman et al., 2019), see Table S1) and cover almost the entire evolutionary history of both animal and plant kingdoms. We performed extensive multi-omics sequence physicochemical analyses concatenating promoter GC-content and TF pIs to evaluate the evolutionary trajectories between promoters and TFs. The results show that the evolutionary trajectories of promoter GC-content and TF pIs share an amazing synchronous increased tune in the animal kingdom, while a synchronous decreased trend in the plant kingdom. The selective pressures of the physicochemical properties drive an evolutionary arms race, namely an antagonistic coevolutionary relationship, between promoters and TFs in the animal kingdom. And the altruistic physicochemical interactions between promoters and TFs engage in a mutualistic coevolution in the plant kingdom. Furthermore, the physicochemical evolution provide additional evidence that Mammalia is closer to Reptilia compared to Aves. These findings provide insights into promoter-TF interactions and biological evolution in molecular interactions and evolution.

## RESULTS

### Physicochemical Specificity of TFs/TF-cofactors and Promoters

TFs and TF-cofactors play different roles in gene expression (Reiter et al., 2017; Thomas and Chiang, 2006). TFs typically bind the TFBSs located in the corresponding promoter to open the double DNA strands of a gene and further control the rate of transcription of its genetic information from DNA to mRNA. TF-cofactors, as intermediary proteins, are recruited by TFs to activate RNA Polymerase II, thereby modulating the expression of genes. Why do they perform differential functions? Note that the spatial conformation of molecules typically affects their functions. The electrical property of amino-acids is one of the most crucial physicochemical properties that shape the specific spatial conformation of proteins. We therefore estimated the electrical property of TFs/TF-cofactors (97 animal species) and whole-proteomes (74 animal species) in terms of pI.

Interestingly, we found that the mean pIs of TFs are significantly higher than those of TF-cofactors at all phylogenetic levels of evolution (Figure 1A, total P = 4.68E-71), and the combinational pIs of TFs and TF-cofactors are also significantly higher (total P = 1.04E-18) than those of the corresponding whole-proteomes (except Arthropoda). Based on the total-mean pI (=6.066) of all whole-proteomes, we concluded that under the same normal physiological condition, the amino-acids with higher pI (>6.066) tend to be positively charged (alkalinity), and those with lower pI (<6.066) appear to be negatively charged (acidity). Thus, TFs/TF-cofactors hold relatively stronger positive charges compared to their corresponding whole-proteome. Note that DNA exhibits one intrinsic negative charge per base at its sugarphosphate backbone (Fritz et al., 2002). It is therefore the case that the TF/TF-cofactor proteins with high pIs preferentially access the strands of DNA.

**Figure 1.**
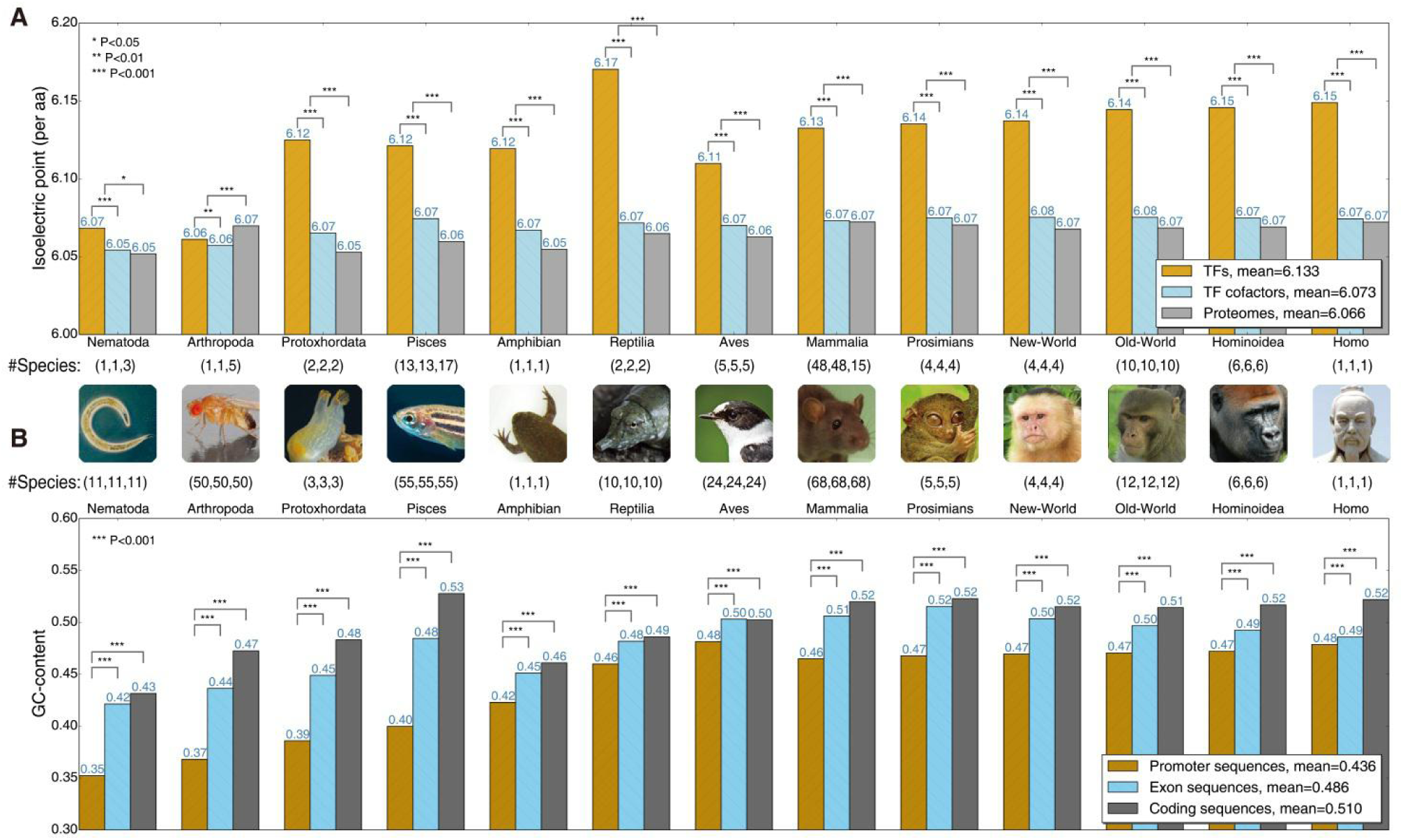
Isoelectric-point (pI) and GC-content Analyses in the Animal Kingdom. (A), (B) The x-axis tick labels show species categories, which follow a phylogenetic relationship (Figure 2B). The animal logos under/over these x-axis tick labels represent their corresponding categories. The elements of each triple like (1,1,3) or (11,11,11) between the x-axis tick labels and the animal logos represents respectively the species numbers of TFs/promoter sequences, TF-cofactors/exon sequences and whole-proteomes/coding sequences. (A) TFs have significantly higher pIs than those of TF-cofactors at all levels of evolution (Table S2). The pIs of the group of TFs and TF-cofactors are also significantly higher than those of the corresponding whole-proteomes (except Arthropoda). (B) In 249 animal species, promoters (8503028 sequences) have significantly lower GC-content than that of exons (49239643 sequences) and coding sequences (7311335 sequences) at all levels of evolution. Together, the pI and GC-content as physicochemical properties can characterize respectively the specificity of protein families and DNA sequences.

We then assessed the GC-content of promoter sequences as well as that of exon and coding sequences in 249 animal species. What we have found is that promoter sequences have significantly lower GC-content than that of exon and coding sequences at all levels of evolution (Figure 1B). It thus suggests that it is easier for promoters to interact with TFs. Taken together, the above findings suggest that pI and GC-content have the potential to represent respectively the specificity of protein families and DNA sequences on physicochemical properties. The TFs/TF-cofactors with higher pIs are more likely to interact with the promoters (TFBSs), especially the lower GC-content promoters.

### Antagonistic Coevolution in the Animal Kingdom

We investigated the evolutionary trajectory of promoter GC-content in 249 animal species (Nematoda to Homo from Ensemble (Hunt et al., 2018)), which are grouped in order by the most probable evolutionary process. The phylogenetic tree of these animals is examplified in Figure 2B by the representative logos of animal categories. What we have found is that the promoter GC-content clearly increases (Figure 2A) in all categories (except Aves). Interestingly, the curves of mammals (Mammalia to Homo) and non-mammals (Nematoda to Aves) share the same monotonic increasing trend (Figure 2A), but the slope of the latter curve is higher, supporting the fact that non-mammals take longer evolutionary years compared to mammals. Following the conventional concept, we use TFs to refer to TFs and TF-cofactors if not explicitly specified in the rest of the paper. We subsequently estimated the changes of pIs of TFs/TF-cofactors in the same evolutionary processes mentioned above, and noticed that the pI values showed an increasing trend based on the pI values (Figure 2C). In particular, the pIs of mammals exhibit a stronger monotonic trend.

**Figure 2.**
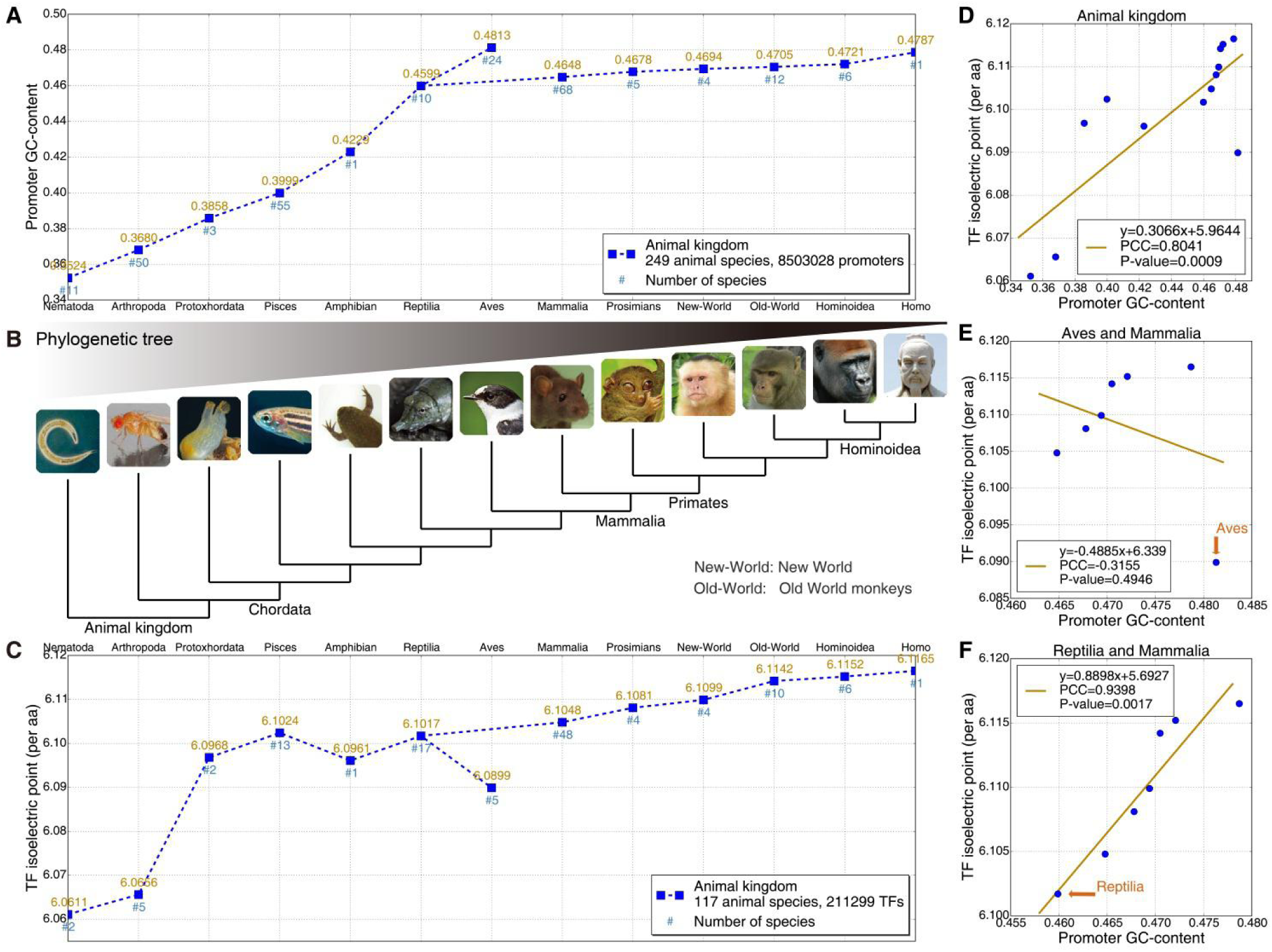
Trends and Correlations of Promoter GC-content and TF Isoelectric-points (pIs) in the Animal Kingdom. (A) and (C) share the same x-axis tick labels, each of which represents an animal species category. The representative animal logos of these categories are listed in order as shown in (B). (A) The promoter GC-content clearly increases in most categories (except Aves). (B) The phylogenetic tree of these animal categories exhibits the (possible) evolutionary process used in (A) and (C) based on Ensemble. (C) The pIs of TFs increase overall. (D) The promoter GC-content and TF pIs satisfy a very strong positive correlation (*PCC* = 0.8041). (E), (F) The location of Aves (outlier) observably departs from those of mammals (*P* = 0.4946), but the point of Reptilia perfectly overlaps with the fitting line (*PCC* = 0.9339), which provides additional evidence that Reptilia is closer to Mammalia compared to Aves. Together, the synchronous increases along with the close correlations of promoter GC-content and TF pIs reveal an antagonistic coevolution between promoters and TFs in the animal kingdom.

Comparing Figure 2A with 2C, we found that they share a similar trend. We therefore explored the relation between promoter GC-content and TF pIs using a scatter diagram. The results suggest that there is a very strong positive correlation between promoter GC-content (Figure 2A) and TF pIs (Figure 2C) in the animal kingdom, with the Pearson-correlation-coefficient (PCC) being 0.8041 and P-value 0.0009 (Figure 2D, using two-tailed t-test) respectively. Close observation of the curves revealed that the point of Aves markedly diverge from the global trajectory in both Figures 2A and 2C. We then analyzed the relation between promoter GC-content and TF pIs from Aves to Homo. The scatter diagram (Figure 2E) shows that Aves is an outlier and its location observably departs from those of mammals (P = 0.4946). Subsequently, we combined Reptilia and Reptilia as a hypothetical evolutionary process and investigated the relation between promoter GC-content and TF pIs. The results (Figure 2F) indicate that the point of Reptilia perfectly overlaps with the fitting line (PCC = 0.9339, P = 0.0017). The regression analysis of promoter GC-content and TF pIs provides additional evidence that Reptilia is closer to Reptilia n lineages compared to Aves. In fact, it has been known that Mammalia evolved from Reptilia (Janes et al., 2010), which in turn supports the correlation between the two physicochemical properties.

In summary, the promoter GC-content and TF pIs follow a mutually strengthening relationship in nearly one billion years of phylogeny. This finding suggests that the close interactions between promoters and TFs in the balance of armament convey an antagonistic coevolution in the animal kingdom. Promoters and TFs affect each other’s evolution under the selective pressures of promoter GC-content and TF electrical property.

### Mutualistic Coevolution in the Plant Kingdom

Does the antagonistic coevolutionary relationship of promoters and TFs exist in the plant kingdom as well? To answer this question, we estimated the relation between promoters and TFs in 223 plant species, which are categorized by the evolutionary process. The phylogenetic tree of these plant categories is shown in Figure 2C. Due to the absence of gymnosperms data in the benchmark databases, the angiosperm species are divided into three sub-groups, lower, medium and higher angiosperms, in order to increase the number of evolutionary categories. By tracking the trend of promoter GC-content in the plant kingdom (62 plant species), we unexpectedly found that the GC-content decreases overall (Figure 3A), which is just opposite to that of the animal kingdom. Estimation of the trend of TF pIs in 161 plant species (Algae to Angiosperm) shows that the pI curve (Figure 3B) decreases overall, which is very similar to the trend of promoter GC-content (Figure 3A). We continued to analyze the scatter diagram (Figure 3D) between promoter CG-content and TF pIs. It is found that they meet a very strong positive correlation (PCC = 0.9333, P = 0.0065). The close synchrony of promoter GC-content and TF pIs suggests that promoters and TFs remain a symbiotic marriage and enjoy a mutualistic coevolutionary relationship under changing promoter GC-content and electrical property in the plant kingdom.

**Figure 3.**
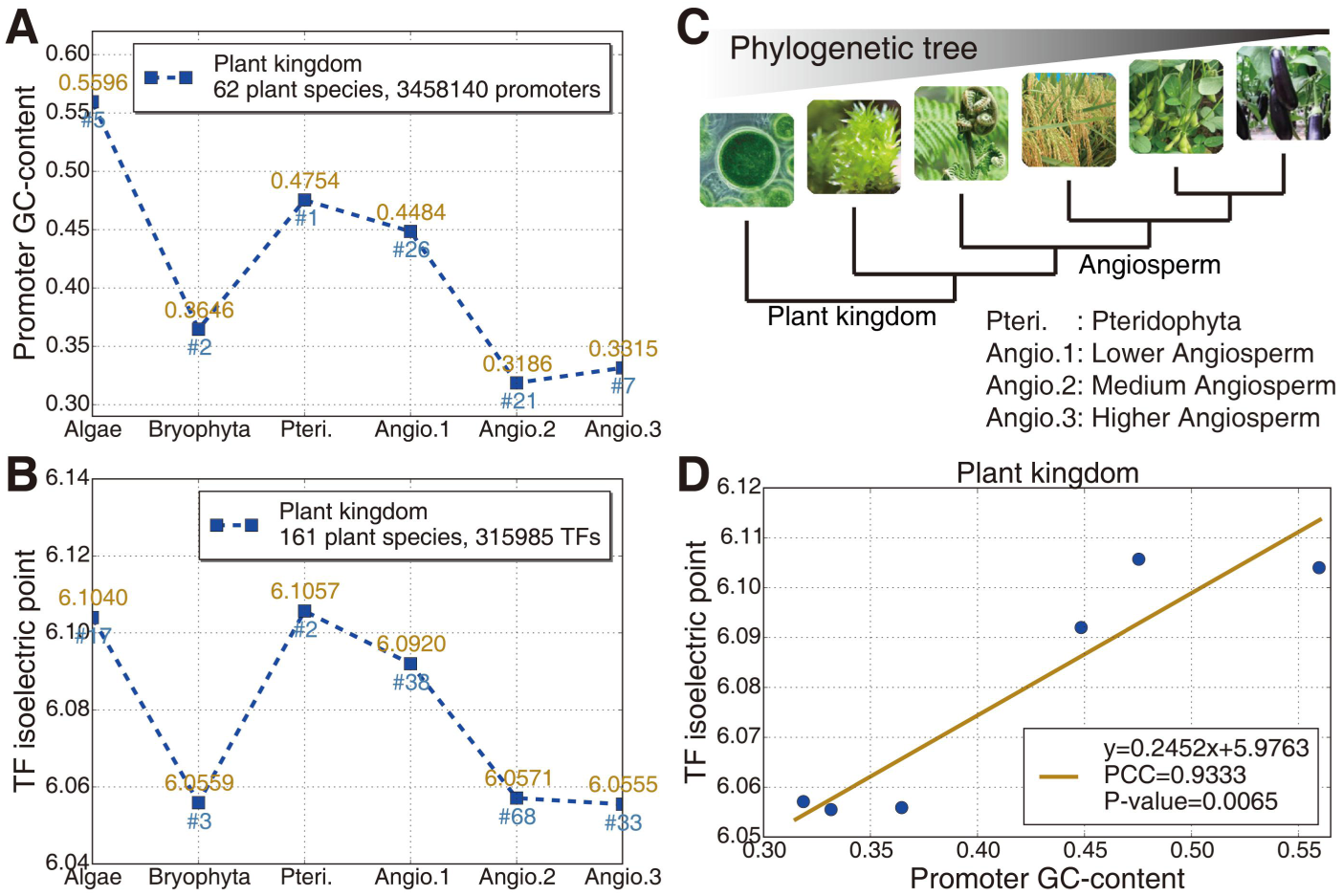
Trends and Correlations of Promoter GC-content and TF Isoelectric-points (pIs) in the Plant Kingdom. (A), (B), (C) The x-axes share the same plant species categories and their representative logos are orderly shown in (C). The numbers on the right of ‘#’ indicate the numbers of species. The phylogenetic tree in (C) of these plant categories indicates the evolutionary process based on EnsemblePlants32. Both the promoter GC-content and TF pIs decrease overall (except Bryophyta). (A) and (B) are characterized by a very similar trend. (D) There is a dramatically positive correlation (*PCC* = 0.9333) between promoter GC-content and TF pIs. Together, the synchrony of promoter GC-content and TF pIs reveals a mutualistic coevolution between promoters and TFs in the plant kingdom.

### Potential Coevolutionary Mechanisms

We next illustrate the coevolutionary strategies and mechanisms between promoters and TFs (or TF complexes) we have unearthed in both animal and plant kingdoms. In the animal kingdom (Figures 4A and 4B), the promoter GC-content clearly increases during the evolutionary process. In this case, the strands of promoter regions tend to be harder to unwind and transcribe, because (1) GC pair has three hydrogen bonds while AT pair has only two bonds; (2) GC-rich regions typically contribute to the base stacking of adjacent bases and therefore block the interactions between promoters and TFs (Yakovchuk et al., 2006). The TF pIs increase following the animal evolution and the strengthened property of positive electricity enables TFs to have more opportunities to trigger the interactions with promoters, as the DNA molecule exhibits one intrinsic negative charge at its backbone of double helix36. The opposite-charges between TFs and DNA molecules drive them to mutually approach each other and closely interact. In summary, while promoters prefer to protect the double-strands from TFs’ unwinding and transcription by increasing the promoter GC-content, so as to maintain the DNA conservation, TFs always strengthen their own capability to bind the TFBSs by increasing the positive electrical property. This observation provides clear evidence for both parasitism and mutualism between promoters and TFs. The selective pressures of the physicochemical properties drive an evolutionary arms race, namely an antagonistic coevolutionary relationship, between promoters and TFs.

**Figure 4.**
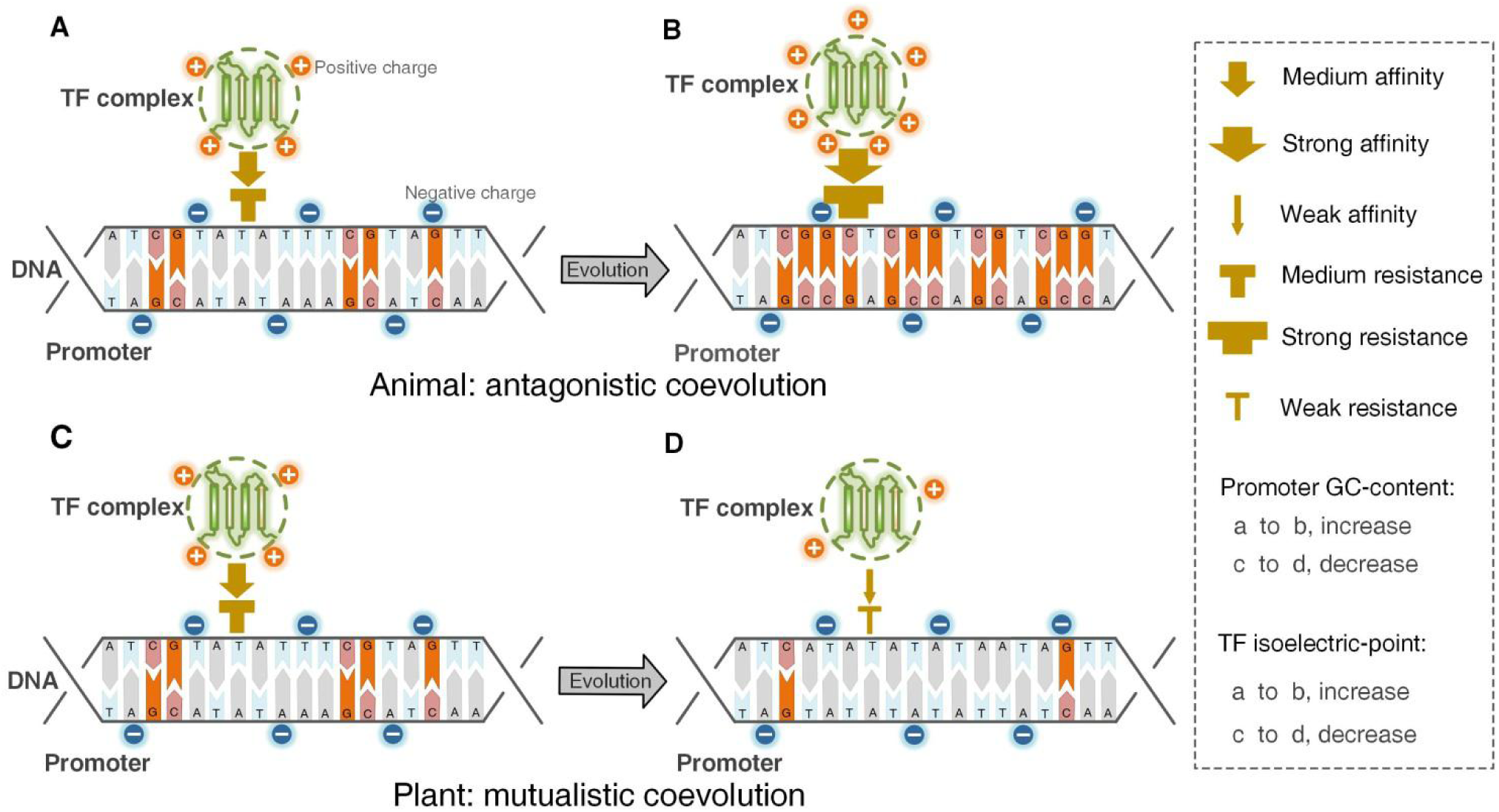
Coevolution of Promoters and TFs (TF complexes) in the Animal and Plant Kingdoms. (A), (B), (C) and (D) represent four different interaction statuses of promoter-TF pairs during the regulation of gene expression in evolutionary processes. The negative charges around the sugarphosphate backbone of promoters are labeled. From (A) to (B), the increase of positive charges carried by TFs reflects that TFs tend to have stronger electrical property and alkalinity. The transition of (A) to (B) features an antagonistic coevolution in the animal kingdom. By contrast, the decrease of positive charges from (C) to (D) renders that the TFs have weaker electrical property and alkalinity. The transition of (C) to (D) conveys a mutualistic coevolution in the plant kingdom.

In the plant kingdom (Figure 4C,D), the promoter GC-content decreases overall, which is contrary to that in the animal kingdom. Such decreased content is more beneficial for performing the functions of TFs during transcription. Interestingly, the positive electrical property of TFs appears to be weaker, which shares a very similar trend with the promoter GC-content. The simultaneous weakening changes of promoter stability and TF activity provide benefits to both partners, thus retaining the symbiosis evolution of molecules in plant cells. Taken together, the altruistic interactions between promoters and TFs engage in a mutualistic coevolution in the plant kingdom.

The evolution of molecular interaction networks drives the diversity of traits and species in biology (Guimaraes et al., 2017; Mann et al., 2016; Paterson et al., 2010) through continual natural selection for adaptation and counter-adaptation. In the animal kingdom, the strengthening of the positive electrical property enhances the DNA binding activity of TFs, and therefore makes more proteins be DNA-binding proteins or TF proteins (Extended Data Figure 1B). The fitting function (y=1.4889x-0.567) of normalized scatter diagram (see STAR Methods) between gene and TF numbers (Figure S1D) suggests that the relative growth rate of TF quantity is nearly 1.5 times faster than that of gene quantity. The increase of TFs produces more powerful TF complexes (gene pleiotropy) and TF-gene links (regulatory links) (Figure 5A to 5B), which contribute to more complex regulatory relationships/networks between promoters and TFs in animals. The promoter-TF antagonistic coevolution is a major catalyst that makes the rapid increase of TFs or regulation links, thus driving the evolution of transcriptional networks between promoters and TFs.

**Figure 5.**
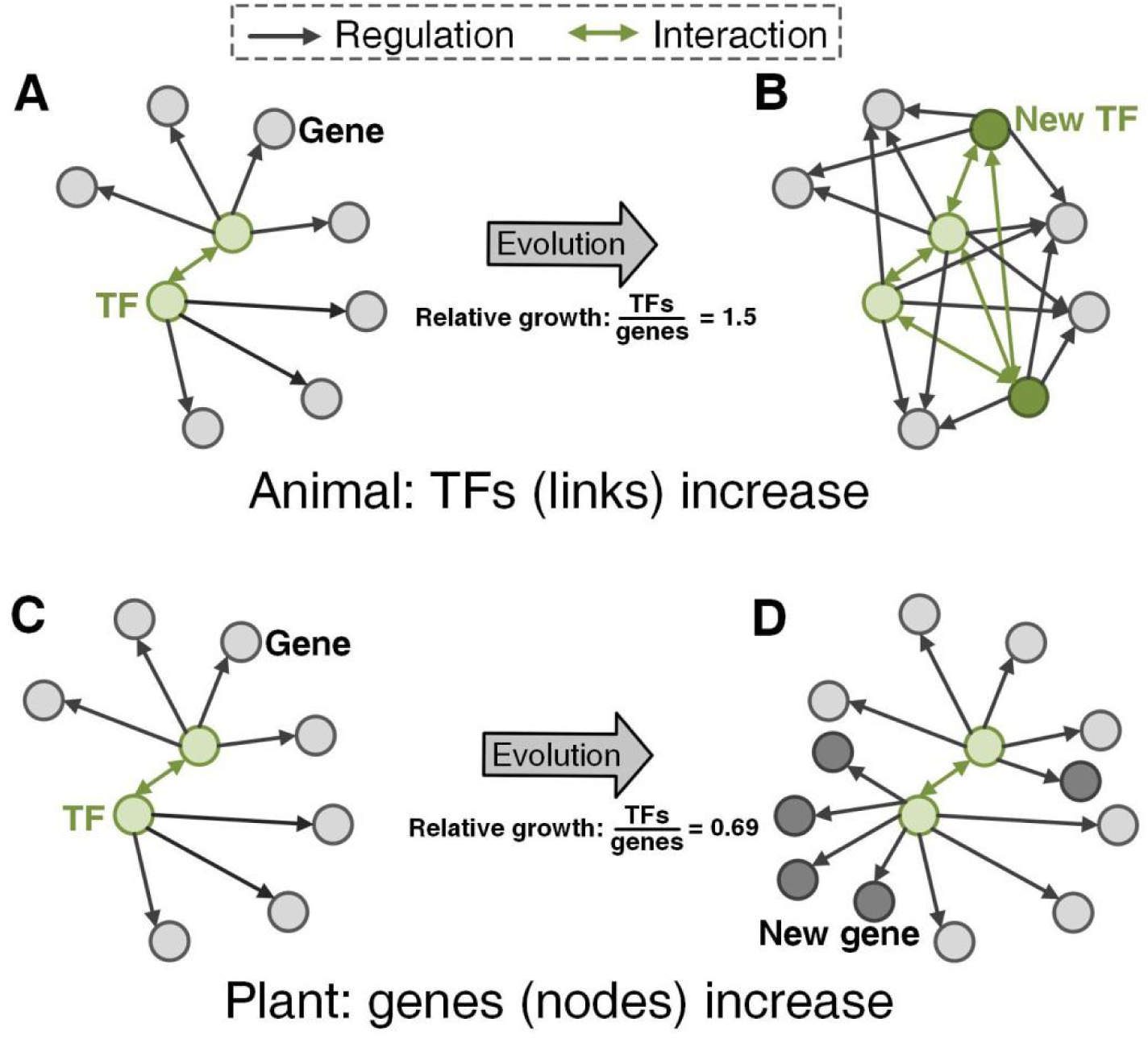
Distinct Gene Regulatory Networks in the Animal and Plant Kingdoms. In animal evolutionary process, the number of TFs increases notably, but that of genes increases dominantly during plant evolutionary process. More precisely, the relative increased speed of TF number is nearly 1.5 times faster than that of gene number in the animal kingdom (Figure S1D), while is nearly 0.69 times slower in the plant kingdom (Figure S2D), which provides another attractive clue for a better understanding of promoter-TF coevolution of animals and plants.

Naturally one may wonder in the plant kingdom, how the mutualistic coevolution between promoters and TFs maintain the evolutionary pace of plants themselves under natural selection or random mutations of DNA? The promoters with low GC-content enable relatively underactive TFs to bind the TFBSs, but may in the present case act for mutual benefit. The fitting function (y=0.6855x-0.0836) of normalized scatter diagram (Extended Data Figure 2D) shows that the relative growth rate of TF quantity is nearly 0.69 times slower than that of gene quantity. The altruistic promoter-TF interactions facilitate the generation of new genes (Extended Data Figure 2A), thereby enhancing the adaptability of plants. The emerging new genes give rise to more complex and diverse transcriptional networks between TFs and promoters/genes (Figure 5C to 5D), which drives the ongoing evolution of plants. In contrast to the increase of regulatory links (new TF-gene links or gene pleiotropy) in the animal kingdom, the species in plant kingdom adopt a different strategy, i.e., increasing regulatory nodes (number of genes or new genes), which drives the evolution of transcriptional networks of species to adapt the changing environment.

The two distinct mechanisms of promoter-TF coevolution in the animal (antagonistic coevolution) and plant (mutualistic coevolution) kingdoms result in different ratios of TF quantity to gene one (Figure S3). The mean ratio in the animal kingdom (0.1039) is significantly (*P* = 0.0078) higher than that in the plant kingdom (0.0608), which supports the evolutionary laws of transcriptional networks in animals and plants. Overall, the evolution of regulatory networks provides further evidence in support of the two distinct coevolutionary mechanisms we have found in the animal and plant kingdoms.

## DISCUSSION

Our physicochemical analyses of massive multi-omics sequences yielded two intriguing findings. First, promoter GC-content accompanying the evolutionary processes exhibits a clear increasing trend in the animal kingdom but a decreasing trend in the plant kingdom. Second, the positive electrical property of TFs is strengthened in the animal kingdom, but is weakened in the plant kingdom. Further examination of these findings by using regression analysis offered the following insights. First, promoters and TFs form a coevolutionary relationship in both animal and plant kingdoms under the selective pressures of promoter GC-content and TF pIs. Second, in the animal kingdom, promoters and TFs show an antagonistic coevolutionary relationship induced by the syntropic changes of promoter GC-content and TF pIs. Third, in the plant kingdom, promoters and TFs show a mutualistic coevolutionary relationship because of the altruistic features of promoter GC-content and positive electrical property of TFs.

Molecular evolution is a fundamental driver of genetic divergence, ontogenesis, and the ongoing trait evolution of species. Because of the vital roles of promoters and TFs in transcriptional regulations of eukaryotic cells, the distinct coevolutionary mechanisms may be the determinants contributing to the genetic divergence between animals and plants from their common ancestors. In particular, the two coevolutionary forms render the transcriptional networks of species in the two kingdoms evolve in different evolutionary transitions, i.e., one (animal kingdom) by increasing network links and the other (plant kingdom) by increasing network nodes. The pervasive antagonistic coevolution is a potentially critical mechanism that drives the animal kingdom (~7.77 million species (Mora et al., 2011; Strain, 2011)) to engender far more diverse species compared to the plant kingdom (~298K species (Mora et al., 2011)) under continual natural selection. Growing evidence shows that both tumors (Francis et al., 2019; Jin et al., 2019; Perera et al., 2016) and ageing (Janes et al., 2018; O’Callaghan and Vassilopoulos, 2017; Stegeman and Weake, 2017) are correlated with the interactions of promoters and TFs. The promoter-TF coevolutionary trajectories in the animal kingdom will shed further light on the mechanisms underlying tumor initiation and progression as well as human ageing, which are yet to be fully understood.

## STAR*METHODS

Detailed methods are provided in the online version of this paper and include the following:

- DATA PREPROCESSING
- GC-CONTENT
- TF AND PROTEOME ISOELECTRIC-POINTS
- NORMALIZED SCATTER DIAGRAM

## SUPPLEMENTAL INFORMATION

Supplemental Information includes three figures and two tables and can be found with this article online at…

## AUTHOR CONTRIBUTIONS

J.S.Z. and L.N.C. designed the study. J.S.Z. and X.T.Y. coded programs and analyzed output data. Z.X.S designed the phylogenetic tree. S.T.H. provided the original datasets. Y.W.Z. performed the statistical analysis. H.D. designed the experiments of isoelectric-points. X.H.H., L.N.C. and J.S.Z. designed the figure of evolutionary mechanisms. K.K.D., H.Q.W, B.H and T.Z. polished the manuscript. L.N.C. and W.W. supervised the experiments. J.S.Z. wrote the manuscript.

## ACKNOWLEDGMENTS

We thank A. Hsueh, L. Tao, J. Guo, and F. Zhang for useful comments on the manuscript. This work is supported by the National Key Research and Development Program of China (2017YFA0505500 to L.N.C., 2017YFC0909502 to X.H.H.); the National Science Foundation of China (61602460 to J.S.Z., 31771476 and 31930022 to L.N.C., 61803360 to X.T.Y., 31771408 to Z.X.S, 11602092 to B.H, 11871456 to T.Z., 11701379 to X.Q.Y.); the Shanghai Municipal Science and Technology Major Project (2017SHZDZX01 to L.N.C.); the Fundamental Research Funds for the Central Universities (3102019JC007 to W.W.); and the Strategic Priority Research Program of CAS (XDB13000000 to W.W.).

