## Supplementary material for "The Coevolution of Promoters and Transcription Factors in Animal and Plant Cells": SI

### STAR★METHODS

#### DATA PREPROCESSING

The datasets including genome and proteome sequences were obtained from several benchmark databases (such as Ensemble, EnsemblePlants, UniProt, AnimalTFDB, and PlantTFDB). Nevertheless, it is inevitable that some sites are equivocal owing to the site mutations or the limitation of sequencing depth. For instance, many sites were labeled as letter N in genome sequences, but labeled as letter X in proteome sequences. Such noises of indeterminate nucleic-acids and amino-acids were taken into consideration in our experiments so as to boost accuracy.

#### GC-CONTENT

GC-content generally is the percentage of guanine or cytosine on a DNA or RNA molecule (Kudla et al., 2006; Smarda et al., 2014; Smith, 2009). In our study, given a promoter sequence, GC-content refers to the sum of the percentages of guanine and cytosine, namely

$$GC\ content = \frac{number\ of\ G + number\ of\ C}{length\ of\ sequence - number\ of\ N} \quad (1)$$

where N denotes the uncertain sites.

##### TF AND PROTEOME ISOELECTRIC-POINTS

Similar to the promoter data, the real TF and proteome sequences also contain much noise. Provided a protein sequence S, the mean isoelectric-point (pI) value is therefore calculated by

$$pI = \frac{\sum_{i=1}^{|S|} I(S_i)}{|S| - |X|} \quad (2)$$

where $\mathrm{I}(S_i)$ represents the pI value of the ith amino-acid in sequence *S* and *X* is the noise. $|S|$ and $|\mathrm{X}|$ are the lengths of sequence S and X respectively.

#### NORMALIZED SCATTER DIAGRAM

Scatter diagram plays an important role in regression analysis (Scharf et al., 1998). However, the general scatter diagram is not enough to reflect the relative changes of two variables by fitting functions. We introduce a modified scatter plot called normalized scatter diagram to characterize the relative variations of increase or decrease. The values of vertical and horizontal axes are limited to [0,1]. It is therefore necessary to normalize each value of both variables for feeding the normalized scatter diagram. The normalized value can be computed by

$$x_i^{'} = \frac{x_i}{\max\limits_{1 \le j \le n} \{x_j\}} \tag{3}$$

where $x_i$ represents a value of the variables, *n* denotes the number of vertical or horizontal variable, and $x_i^{'} \in [0,1]$ is the normalized value of $x_i$. For the relative positions of variable points and the fitting trend-lines, there is no difference between the general scatter diagram and the normalized one, while the fitting functions of them differ from each other. Compared to the general scatter diagram (Figures S1C and S2C), the slope of fitting function in normalized scatter diagram better indicates the relative changes of two variables (Figures S1D and S2D).

#### Supplemental Figures

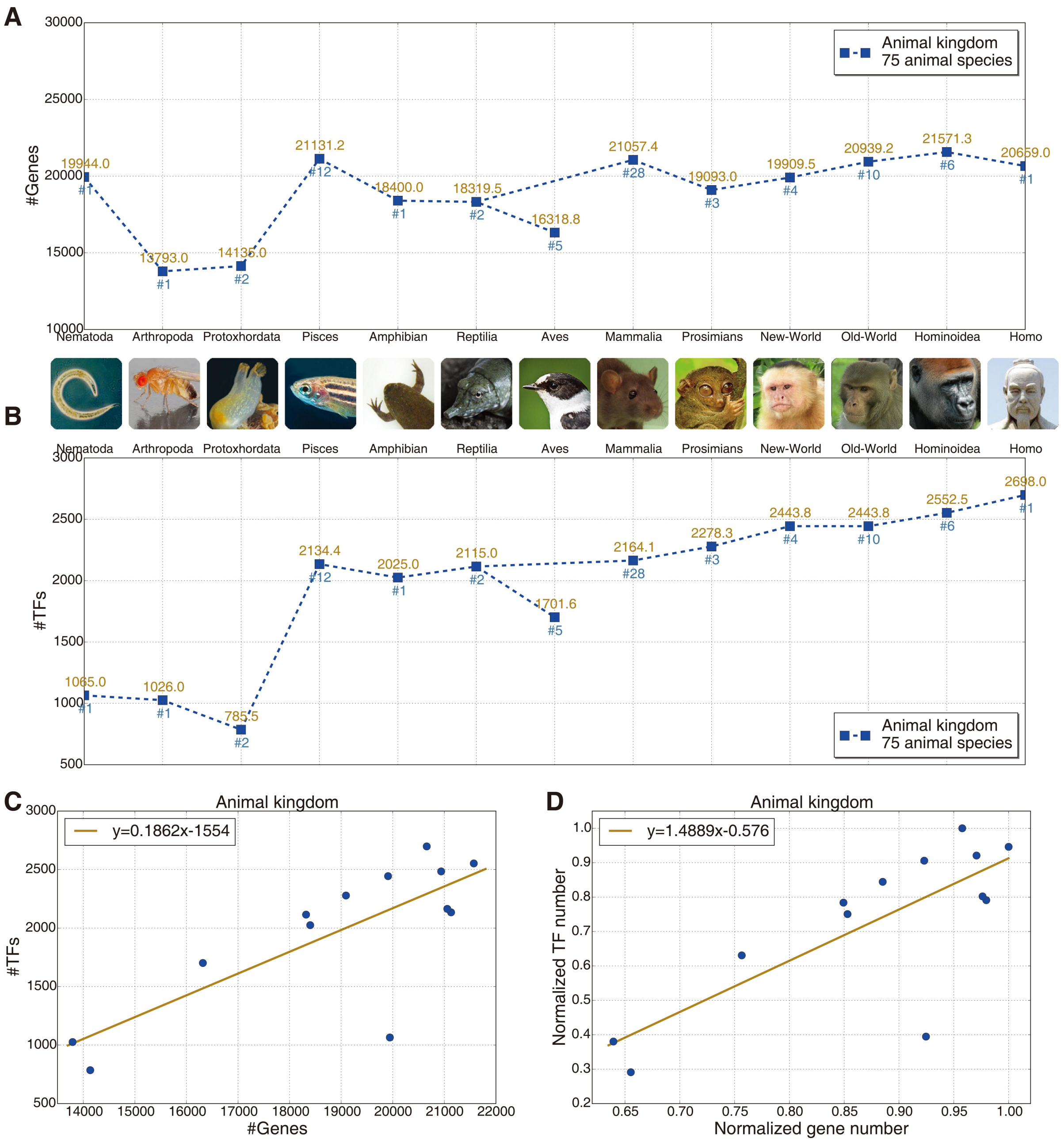


**Figure S1. Tends of Gene and TF Numbers in the Animal Kingdom.**

(A) and (B) share the same x-axis tick labels, each of which represents an animal species category. The numbers on the right of '#' indicate the numbers of species. The representative animal logos of these animal categories are listed between (A) and (B). The phylogenetic tree of the animal categories is shown in Figure 2B. The mean numbers of animal genes slightly increase overall, while those of animal TFs show a more salient growth. (C) reports the scatter diagram of gene and TF numbers. (D) The fitting function (*y=1.4889x-0.567*) of normalized scatter diagram suggests that TFs increase dominantly (1.4889>1) compared to genes in the animal kingdom.

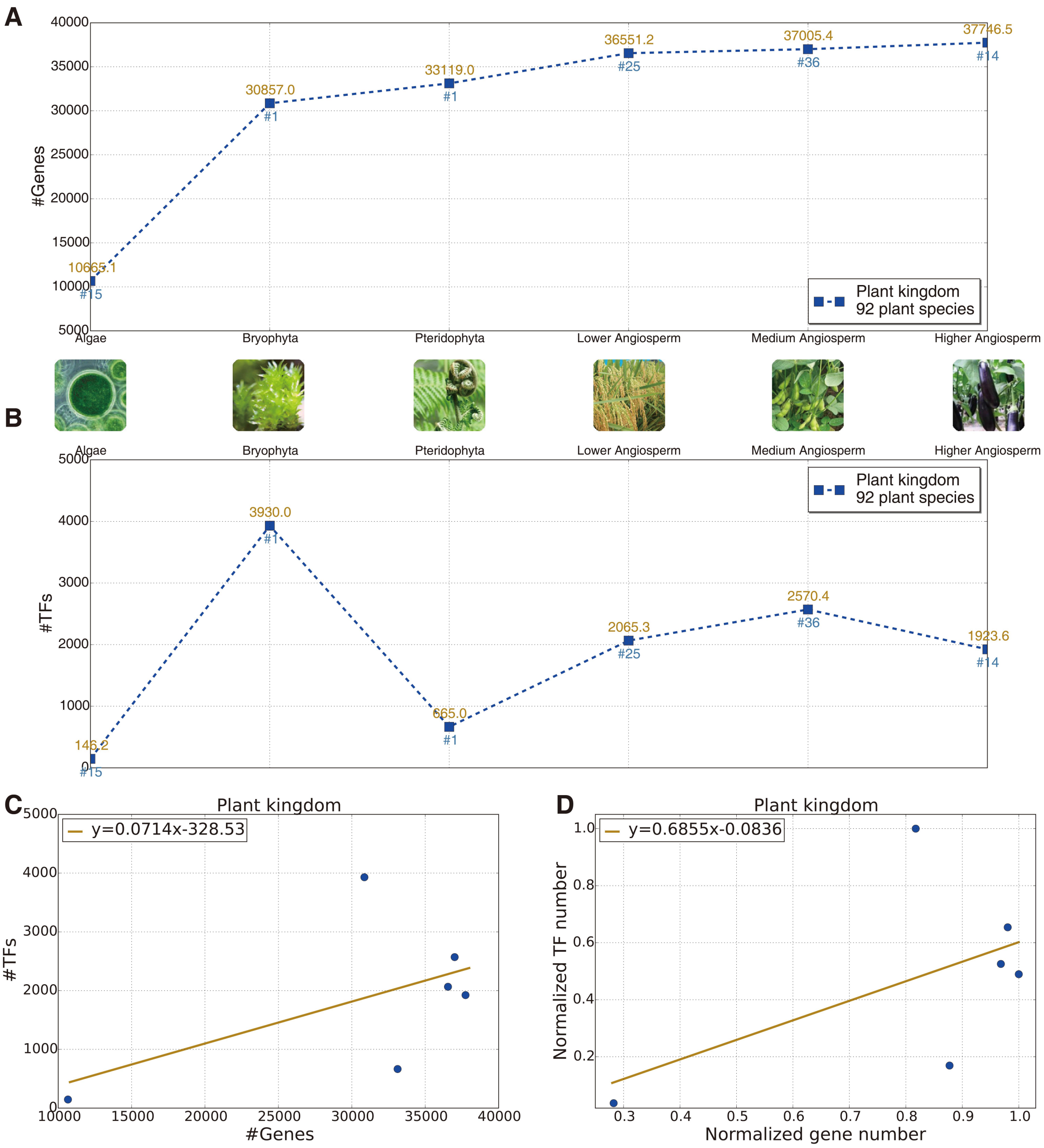


**Figure S2. Tends of Gene and TF Numbers in the Plant Kingdom.**

(A) and (B) share the same x-axis tick labels, each of which represents an animal species category. The numbers on the right of '#' indicate the numbers of species. The representative animal logos of these categories are listed between (a) and (b). The phylogenetic tree of the animal categories is shown in Figure 3C. The mean numbers of plant genes increase dramatically. By contrast, the mean numbers of plant TFs show a slightly increased tend overall. (C) reports the scatter diagram of gene and TF numbers. (D) The fitting function (*y=0.6855x-0.0836*) of normalized scatter diagram suggests that genes increase dominantly (0.6855<1) compared to TFs in the plant kingdom.

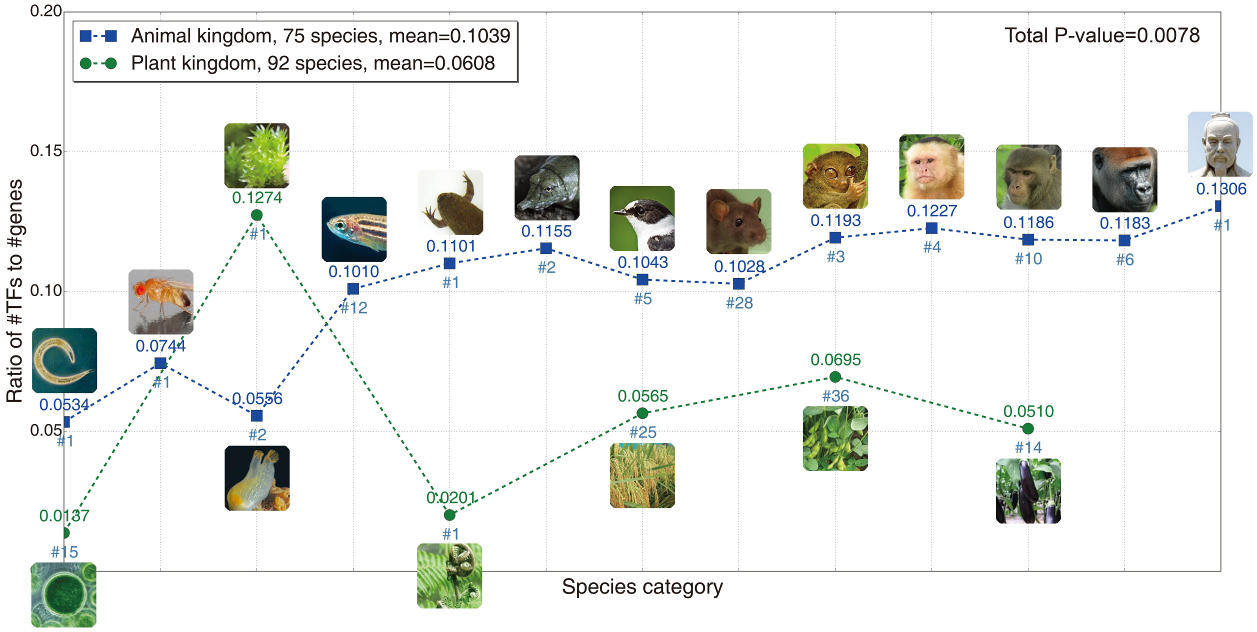


**Figure S3. Ratios of TFs Numbers to Gene Numbers in the Animal and Plant Kingdoms.**

The animal or plant logos represent species categories, all of which are listed near the animal or plant curves. The numbers on the right of '#' indicate the numbers of species. The mean ratio of TFs numbers to gene numbers in the animal kingdom (0.1039) is significantly (*P*= 0.0078) higher than that in the plant kingdom (0.0608), which further supports the two distinct evolutionary mechanisms we have found in animals and plants.

### Supplemental Tables

**Table S1. Dataset Summary.** The datasets involved in our study include both genome and proteome sequences and cover almost the entire evolutionary history of both animal and plant kingdoms.

| For | Dataset name | #Species | #Sequences | Database |
|---|---|---|---|---|
| Fig.1 | Animal TFs | 97 | 125145 | AnimalTFDB |
| | Animal TF-cofactors | 97 | 80058 | AnimalTFDB |
| | Animal proteomes | 74 | 2132385 | Ensemble, UniProt |
| | Animal promoters | 249 | 8503028 | Ensemble |
| | Animal exons | 249 | 49639643 | Ensemble |
| | Animal coding sequences | 249 | 7311335 | Ensemble |
| Fig. 2 | Animal promoters | 249 | 8503028 | Ensemble |
| | Animal TFs/TF-cofactors | 117 | 211299 | AnimalTFDB, Ensemble, Uniprot |
| Fig. 3 | Plant promoters | 62 | 3458140 | EnsemblePlants |
| | Plant TFs/TF-cofactors | 161 | 315985 | PlantTFDB |

**Table S2. Isoelectric-point (pI) T-test (P-value) of TFs/TF-cofactors and Proteomes.** The second column shows the P-value of T-Test between TFs and TF-cofactors, and the third one between TFs/TF-cofactors and whole-proteomes.

| Categories | TFs & TF-cofactors | TFs/TF-cofactors & whole-proteomes |
|---|---|---|
| Nematoda | 4.02E-05 | 0.0492 |
| Arthropoda | 1.66E-03 | 5.08E-06 |
| Protoxhordata | 1.90E-21 | 1.05E-14 |
| Pisces | 0 | 0 |
| Amphibian | 2.77E-24 | 1.55E-27 |
| Reptilia | 8.43E-112 | 3.36E-142 |
| Aves | 1.88E-138 | 6.45E-67 |
| Mammalia | 0 | 0 |
| Prosimians | 4.87E-131 | 2.41E-127 |
| New World monkeys | 4.47E-134 | 2.38E-161 |
| Old World monkeys | 0 | 0 |
| Hominoidea | 3.35E-226 | 1.91E-269 |
| Homo | 2.97E-41 | 8.05E-35 |
